# BRISANCE: A GPU-Accelerated Pipeline for Somatic Short-Variant Detection in Whole-Genome Sequencing Data

**DOI:** 10.64898/2026.11.24.768872

**Authors:** Murilo Porfírio de Aguiar, Saloe Bispo, Silvia Aparecida Teixeira, Rui Manuel Reis, Luciane Sussuchi da Silva, Letícia Graziela Costa Santos

## Abstract

Post-sequencing computational processing of tumor–normal whole-genome sequencing (WGS) remains computationally intensive and can delay somatic variant detection in clinical and translational workflows. We developed and evaluated BRISANCE, a graphics processing unit (GPU)–accelerated pipeline based on NVIDIA Parabricks for detecting somatic single-nucleotide variants and short insertions/deletions, using matched tumor–normal WGS data from the Cancer Genome in a Bottle HG008 pancreatic ductal adenocarcinoma model. Concordance with the DRAGEN-derived comparator call set was assessed using allele-level variant-set comparison, RTG Tools haplotype-aware benchmarking, and IGV manual review of four literature-reported HG008-T sample loci. Runtime was measured in two analyses: a fixed CPU–GPU comparison and GPU-scaling experiments. At the analyzed whole-genome call-set level, the PASS-filtered, positive-TLOD Parabricks set and the PASS-filtered DRAGEN set contained 17,134 and 16,885 variants, respectively, sharing 14,430 variants and yielding an intersection-over-union overlap of 73.66%. Shared fractions were 76.09% for single-nucleotide variants and 69.06% for insertions/deletions. Using the DRAGEN-derived call set as a comparator rather than a validated truth set, RTG score-based filtering increased comparator-based precision from 0.8375 to 0.9110 and F-measure from 0.8569 to 0.8821. Qualitative IGV review showed consistent read-level support at the four selected HG008-T loci. In the fixed CPU–GPU comparison using the HG008 tumor–normal dataset, the summed runtime of the evaluated preprocessing and somatic variant-calling components was 10.6 hours for the GPU-accelerated workflow versus 219.4 hours for the CPU-based workflow, corresponding to a 95.1% reduction and a 20.6-fold speedup under the evaluated conditions. In GPU-scaling experiments, alignment/preprocessing reached its shortest runtimes with 6–8 GPUs, whereas MutectCaller reached its shortest runtime with 2–3 GPUs, depending on CPU-thread configuration. Full-input Nextflow execution reproduced the historical CHROM/POS/REF/ALT record set, and a configuration-based interface was developed for user-defined analyses.

**Highlights:**

- The tested GPU workflow achieved a 20.6-fold speedup over the CPU workflow in HG008.
- RTG filtering increased F-measure against DRAGEN from 0.8569 to 0.8821.
- Shortest runtimes differed by workflow step, GPU count, and CPU-thread setting.
- GPU acceleration enables substantially faster bioinformatics workflows while supporting step-specific optimization of computational resources.

## INTRODUCTION

Cancer develops through the progressive accumulation of somatic genomic alterations that disrupt core biological processes, including cell proliferation, survival, differentiation, DNA repair, and genome maintenance (1,2). Although tumors may harbor thousands of somatic mutations, only a limited subset contributes directly to tumor initiation, progression, or therapeutic response and is therefore classified as driver alterations; the remaining changes are generally considered passenger events (1,3). Large-scale sequencing studies have demonstrated that the number, type, and genomic distribution of these alterations vary substantially across cancer types and individual tumors, reflecting marked molecular heterogeneity (3,4). Accurate identification of somatic variants is therefore essential for characterizing tumor biology and supporting genomic analyses in research and precision oncology (5,6).

Molecular profiling is now integral to precision oncology because tumor genomic alterations can guide diagnosis, prognosis, and treatment selection across cancer types (5,6). Examples include *EGFR* mutations and *ALK* rearrangements in non-small cell lung cancer, *ERBB2/HER2* alterations in breast and other solid tumors, agnostic markers such as *BRAF* mutations and *NTRK* fusions, and *BRCA1/2* alterations as biomarkers for DNA damage–targeted therapies (7–13). Professional society recommendations support routine next-generation sequencing in selected metastatic cancers (14).

Next-generation sequencing is a principal method for detecting cancer-associated genomic alterations because it analyzes many genomic loci in parallel (14). Sequencing instruments generate raw read data, typically in FASTQ format, rather than clinically interpretable variant calls in a human readable format (15). Before biological interpretation or clinical reporting, these data require a processing workflow that includes alignment to a reference genome, duplicate marking, base quality score recalibration, somatic variant calling, and downstream filtering to produce analysis-ready files and confident variant call sets (15,16).

Converting raw sequencing reads into analysis-ready files and somatic variant calls remains a practical bottleneck in genomic oncology because routine workflows can exceed two weeks in real-world settings, limiting use in time-sensitive clinical and translational scenarios (17). Graphics processing unit (GPU)–based computational strategies can reduce genomic workflow runtime while preserving compatibility with established variant-calling frameworks (18). A successful example of a GPU–based computational strategy is the NVIDIA Clara Parabricks software suite, which provides GPU-accelerated implementations of germline and somatic genomic workflows, including Mutect2-based analyses (18,19). Here, we implemented and evaluated a GPU-accelerated NVIDIA Clara Parabricks pipeline for somatic short-variant detection using matched tumor–normal whole-genome sequencing data from the Cancer Genome in a Bottle HG008 pancreatic ductal adenocarcinoma model (18,20). We assessed variant-call concordance against a DRAGEN-derived comparator set using allele-level comparison, haplotype-aware benchmarking with RTG Tools, and manual IGV review, and we evaluated computational performance through a fixed CPU–GPU comparison and GPU-scaling experiments. To facilitate reuse of the pipeline, we also implemented BRISANCE as a Nextflow workflow and developed a configuration-based interface that allows users to specify their paired-end tumor–normal input files, reference resources, and computational settings before automated execution.

## METHODS

### EVALUATION DATASET

Pipeline evaluation was performed using matched tumor–normal whole-genome sequencing data from one individual, HG008, provided by Cancer Genome in a Bottle (Cancer GIAB), comprising a pancreatic ductal adenocarcinoma-derived tumor cell line and matched normal material (21). The Cancer GIAB consortium provides raw sequencing reads in FASTQ format and also distributes a somatic “hard-filtered” VCF generated with Illumina’s DRAGEN pipeline, which was used here as a baseline comparator (22). All files were stored on an on-premises NVIDIA DGX A100 system, and file integrity was verified using MD5 checksums provided by Cancer GIAB (23).

### GPU-ACCELERATED PIPELINE DESIGN

A GPU-accelerated tumor–normal pipeline was built following best-practice guidelines for short-variant discovery (SNVs and short indels) (24) and executed in Docker version 27.1.2 (25). The workflow used NVIDIA Parabricks version 4.5.1-1 (container image nvcr.io/nvidia/clara/clara-parabricks:4.5.1-1) for read alignment and preprocessing (sorting, duplicate marking, BQSR, and indexing), followed by somatic SNV/indel calling. HG008 WGS FASTQ files were processed with NVIDIA Clara Parabricks fq2bam function using the GIAB-provided GRCh38/hg38 reference genome and known-sites VCF files (Mills/1000G indels, known indels, and high-confidence SNPs) for BQSR. Tumor–normal calling was performed with MutectCaller (Parabricks’ accelerated implementation version), producing a tumor–normal somatic VCF. The BRISANCE source code and execution documentation, including the Nextflow workflow and user configuration interface, are available in the project repository (26).

### VARIANT-SET COMPARISON BETWEEN PIPELINES

To compare PASS-filtered variant sets between the GPU pipeline and the GIAB-provided DRAGEN hard-filtered baseline, the raw Parabricks MutectCaller VCF was first processed with GATK FilterMutectCalls using the corresponding MutectCaller statistics file and the GRCh38 reference genome. Variants with FILTER=PASS were first selected using BCFtools view. For the comparison reported here, the Parabricks call set was further restricted to records with a non-missing positive INFO/TLOD value (TLOD > 0) and compared with PASS records from the baseline VCF dragen_4.2.4_HG008-mosaic_tumor.hard-filtered.vcf.gz (22). Comparative summary statistics and distribution-based outputs were first generated with BCFtools stats (Docker image staphb/bcftools:1.24) (27). Final shared and pipeline-specific counts for SNVs and indels were then obtained using a custom Python script for post-processing of the two VCF files. Variants were considered matching when they shared the same chromosome, genomic position, reference and alternate alleles (CHROM, POS, REF, ALT). To assess whether allele fraction contributed to SNV discordance, tumor-sample FORMAT/AF values were extracted using BCFtools query for shared and pipeline-specific SNVs and summarized in four descriptive intervals: <2%, 2% to <5%, 5% to <10%, and ≥10%. For pipeline-specific SNVs, BCFtools isec was additionally used to determine whether the exact allele was absent from the complete alternative-pipeline VCF or present with a non-PASS classification, in which case the corresponding FILTER tags were summarized. These VAF intervals were used only for descriptive stratification and were not applied as additional variant-calling filters. Symbolic alleles were excluded from the custom comparison step. Indels were compared using the same allele-level matching rule, and insertions and deletions were additionally summarized separately. The variant-set comparison scripts and documented analysis procedures are available in the BRISANCE repository (26).

### IGV-BASED QUALITATIVE REVIEW OF SELECTED HG008-T VARIANTS

Selected HG008-T somatic variants reported by McDaniel et al. (2025) were manually reviewed in IGV in both the GIAB-provided BWA-MEM tumor alignment and the Parabricks-derived fq2bam tumor alignment (20). The loci were selected based on their identification in the original HG008 characterization as potentially relevant somatic alterations, including variants in cancer-associated genes and representatives of both SNVs and a short insertion. The reviewed loci were *KRAS* c.35G>T, *TP53* c.395A>C, *INSYN1* c.457C>T, and *SMAD4* c.153dup. For each site, read-level support for the alternate allele, alignment patterns, and the apparent VAF estimated from the IGV nucleotide counts were compared between alignments.

### BENCHMARKING AGAINST A DRAGEN-DERIVED BASELINE SET USING RTG TOOLS

Variant-calling performance was assessed against the Cancer GIAB DRAGEN hard-filtered somatic VCF, which was used as the baseline comparator (22). True-positive, false-positive, and false-negative classifications and the resulting precision, sensitivity, and F-measure values were defined relative to the DRAGEN comparator and should therefore be interpreted as measures of agreement with that call set rather than independently validated analytical accuracy. The GRCh38 reference FASTA used for read alignment (GRCh38_GIABv3_no_alt_analysis_set_maskedGRC_decoys_MAP2K3_KMT2C_KCNJ18. fasta) was converted to RTG Sequence Data Format (SDF) using rtg format within the Docker image realtimegenomics/rtg-tools:3.12.1 (28), producing the reference directory ref_sdf_rtg (SDF version 13). Haplotype-aware comparison between the PASS-filtered baseline VCF and the PASS-filtered, positive-TLOD GPU-derived call set was performed using RTG Tools vcfeval in annotate mode with the Docker image ghcr.io/uclahs-cds/rtg-tools:dev (29). Sample matching was explicitly specified as HG008-T-mosaic, somatic, and benchmarking was run with --squash-ploidy, --ref-overlap, and --threads 8 parameters. For score-based analyses, the INFO.RTG_SCORE field was added to the calls VCF using the caller-specific confidence metric (e.g., TLOD) and provided to RTG via --vcf-score-field INFO.RTG_SCORE. The annotated outputs (baseline.vcf.gz and calls.vcf.gz) were then processed with rtg vcf2rocplot using -f INFO.RTG_SCORE, and ROC-style and precision–recall-like curves were generated from weighted_roc.tsv.gz with rtg rocplot. All RTG analyses were performed in Docker containers. Code and implementation details for reproducing this benchmarking workflow are provided in the Availability of data and materials section.

### CPU PIPELINE FOR RUNTIME COMPARISON

A CPU-based pipeline was executed using containerized tools representative of a conventional best-practices workflow to provide a runtime comparator (30). Read alignment was performed with BWA-MEM version 0.7.19-r1273 using the staphb/bwa:latest Docker image (31), followed by sorting and indexing with SAMtools version 1.21 using the staphb/samtools:latest Docker image (32). Explicit CPU (--cpus=48) and memory (--memory=128g) limits were applied to ensure stable resource usage. Duplicate reads were marked using MarkDuplicates from GATK version 4.6.2.0. Base-quality score recalibration tables were subsequently generated and applied using BaseRecalibrator and ApplyBQSR, respectively, from the same GATK version (33,34), with the same known-sites resources as in the GPU workflow. Somatic variant calling was then performed using Mutect2 from GATK version 4.6.2.0 within the broadinstitute/gatk:latest Docker image (35).

### RUNTIME MEASUREMENT AND PARALLEL SCALING

Elapsed wall-clock time was recorded for each major workflow step and analyzed in two performance comparisons. First, summed preprocessing and somatic variant-calling runtimes were compared using a fixed NVIDIA Parabricks v4.5.1-1 configuration and a conventional CPU workflow following GATK Best Practices–style processing (36) to estimate the overall runtime reduction associated with GPU acceleration (26). Second, intra-GPU scalability was evaluated by varying GPU count and CPU-thread parameters across selected workflow components; these scaling runs were analyzed separately from the fixed CPU–GPU comparison. The execution parameters used for the fixed CPU–GPU comparison is summarized in **Table 1**.

**Table 1.** Execution parameters used for the fixed CPU–GPU runtime comparison.

| **Workflow** | **Step** | **Execution configuration** |
|---|---|---|
| CPU-based | BWA-MEM alignment | 32 threads |
|  | samtools sorting/indexing | 16 threads |
|  | GATK MarkDuplicates | up to 48 CPUs and 128 GiB RAM |
|  | GATK BaseRecalibrator | up to 32 CPUs and 64 GiB RAM |
|  | GATK ApplyBQSR | up to 48 CPUs and 128 GiB RAM |
|  | GATK Mutect2 | 32 threads |
| GPU-accelerated | Parabricks fq2bam | 1 GPU and 16 threads |
|  | Parabricks ApplyBQSR | 1 GPU and 16 worker threads |
|  | Parabricks MutectCaller | 1 GPU and 32 threads |

CPU-based steps were executed using docker containerized tools, except where otherwise specified. In the GPU-accelerated workflow, fq2bam integrates alignment, sorting, duplicate marking, BQSR table generation, and indexing. Both workflows used the same GRCh38 reference genome file and the same known-sites resources for base quality score recalibration. Runtime measurements focused on workflow components used for performance evaluation, whereas read quality control and trimming steps were performed separately and were not included in the metrics.

For the GPU-scaling analysis, runtime was quantified separately for somatic alignment, germline alignment, and variant calling. The accelerated workflow was implemented using NVIDIA Clara Parabricks version 4.5.1-1 (37) and included alignment and preprocessing with fq2bam (38), together with somatic variant-calling workflows based on MutectCaller (39) and DeepSomatic (40). MutectCaller analyses used reheadered BAM files together with recalibration tables generated during preprocessing, whereas DeepSomatic analyses were performed using the tumor and normal BAM files without incorporating recalibration tables. For each GPU-scaling test, the number of GPUs assigned to the job varied from 1 to 8. CPU-thread parameters were tested at 16, 24, and 32 threads. These values refer to the thread parameter passed to the corresponding Parabricks command and should not be interpreted as the final total number of host CPU threads requested by the application.

Thread-related parameters in Parabricks are applied on a per-GPU or per-stream basis depending on the tool. For fq2bam and MutectCaller, the effective thread allocation scales with the number of GPUs (38,39,41), approximately as:

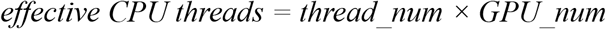

For DeepSomatic, the effective CPU-thread allocation additionally depends on the number of streams, approximately as (40,41):

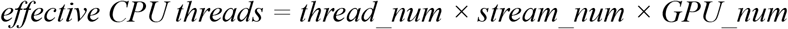

### COMPUTE ENVIRONMENT

All analyses were executed on an on-premises NVIDIA DGX A100 system running Ubuntu 22.04.5 LTS (Jammy Jellyfish) with Linux kernel 5.15.0-1071-nvidia. The system software stack corresponded to NVIDIA DGX Server for DGX A100 (DGX OTA version 6.3.1; DGX SWBUILD version 5.5.0), updated to OTA 6.3.1 on 29 November 2024. The host comprised two AMD EPYC 7742 processors (64 cores per socket; 256 logical CPUs total), 2.0 TiB of system memory, and eight NVIDIA A100-SXM4 GPUs with 80 GB HBM2 memory each (compute capability 8.0), using NVIDIA driver 535.216.03 with CUDA driver/runtime version 12.2. Containerized execution was performed with Docker Engine 27.1.2, with NVIDIA Container Toolkit 1.17.4 and nvidia-docker2 2.13.0-1 enabling GPU passthrough. The host CUDA compilation toolchain corresponded to CUDA toolkit 11.5 (nvcc 11.5.119).

### NEXTFLOW IMPLEMENTATION AND FUNCTIONAL EVALUATION

BRISANCE was implemented as a modular Nextflow DSL2 workflow. The HG008 evaluation used Nextflow version 26.04.6 (42), Java version 21.0.10 (43), and Docker Engine version 27.1.2 (25) on the DGX A100 system described above. Separate processes performed paired-end read trimming and quality control, alignment and BQSR table generation, BQSR application, read-group adjustment, tumor–normal somatic variant calling, VCF compression and indexing, variant filtering, PASS selection, VEP annotation, and positive-TLOD selection. Input samples and FASTQ pairs were specified in a CSV samplesheet, and process dependencies connected the outputs of each stage to the corresponding downstream inputs. Execution generated software-version records, configuration records, task-level execution traces, and HTML execution reports and timelines.

Functional evaluation was performed in a single integrated run using original HG008 reads. Inputs comprised four paired tumor FASTQ inputs and one paired normal FASTQ input, totaling ten compressed files and 500.24 GB. Agreement with the historical output generated outside Nextflow was assessed by exact matching of VCF records using chromosome, position, reference allele, and alternate allele (CHROM, POS, REF, ALT).

Based on the GPU-scaling results, step-specific configurations were selected to balance processing time and computational resource demand. Alignment with fq2bam used three GPUs, a BWA CPU thread-pool parameter of 16 per GPU, and container limits of 64 logical CPUs and 256 GB RAM, with a sorting and postsorting memory limit of 128 GB. MutectCaller used two GPUs, with num-htvc-threads set to 24 and container limits of 64 logical CPUs and 256 GB RAM. Separate ApplyBQSR processing used one GPU, a worker-thread parameter of 16, and container limits of 24 logical CPUs and 128 GB RAM. The two sample alignments were executed sequentially to avoid concurrent three-GPU jobs.

### USER CONFIGURATION AND EXECUTION INTERFACE

A configuration-based interface was developed to support user-defined analyses. An editable INI file specifies paired-end tumor and normal FASTQs, reference and known-sites resources, the VEP cache, trimming settings, GPU devices, and CPU, memory, and thread limits. Before execution, the interface displays an analysis summary and storage estimate for user approval. It then checks software dependencies, container images, and NVIDIA GPU access, offering any required image downloads with explicit consent. A final confirmation is requested before launching the workflow, and the configuration and execution reports are retained with the analysis.

## RESULTS

### VARIANT-SET COMPARISON BETWEEN PIPELINES

At the analyzed call-set level, the PASS-filtered, positive-TLOD Parabricks set contained 17,134 SNVs and indels, whereas the PASS-filtered GIAB-provided DRAGEN baseline contained 16,885. Of these, 14,430 were shared between pipelines, while 2,704 were exclusive to Parabricks and 2,455 were exclusive to DRAGEN, corresponding to an overall overlap of 73.66% across the combined SNV/indel set.

For SNVs, 9,760 calls were shared between pipelines, with 1,609 unique to Parabricks and 1,458 unique to DRAGEN, corresponding to a shared fraction of 76.09%. Low-VAF SNVs were more frequent among pipeline-specific calls than among shared calls (23.2% versus 0.64% for Parabricks and 17.9% versus 1.36% for DRAGEN). Nevertheless, more than three quarters of pipeline-specific SNVs had VAFs of at least 5%, indicating that low VAF characterized only a subset of the discordant calls. Most pipeline-specific SNVs were absent from the complete alternative-pipeline VCF (88.6% of Parabricks-only and 76.7% of DRAGEN-only calls); among cross-detected non-PASS calls, mapping-quality and weak-evidence filters predominated in DRAGEN, whereas strand bias predominated in Parabricks **(Supplementary Table S1)**. For indels, 4,670 calls were shared, while 1,095 and 997 were exclusive to Parabricks and DRAGEN, respectively, corresponding to an overlap of 69.06%. Concordance was therefore higher for SNVs than for indels **(Table 2)**.

**Table 2.** Comparison of SNV and indel call sets between the PASS-filtered, positive-TLOD Parabricks call set and the PASS-filtered GIAB-provided DRAGEN baseline.

| **Category** | **Parabricks** | **DRAGEN** | **Shared (%)** | **Parabricks only** | **DRAGEN only** |
|---|---|---|---|---|---|
| SNVs | 11,369 | 11,218 | 9,760 (76.09%) | 1,609 | 1,458 |
| InDels | 5,765 | 5,667 | 4,670 (69.06%) | 1,095 | 997 |
| Total (SNVs + InDels) | 17,134 | 16,885 | 14,430 (73.66 %) | 2,704 | 2,455 |

### IGV-BASED QUALITATIVE REVIEW OF SELECTED HG008-T VARIANTS

IGV-based visual inspection of selected HG008-T variants showed concordant read-level support between the GIAB-provided BWA-MEM alignment **(Figure 1A)** and the Parabricks-derived fq2bam alignment **(Figure 1B)** across all four literature-reported loci.

**Figure 1.**
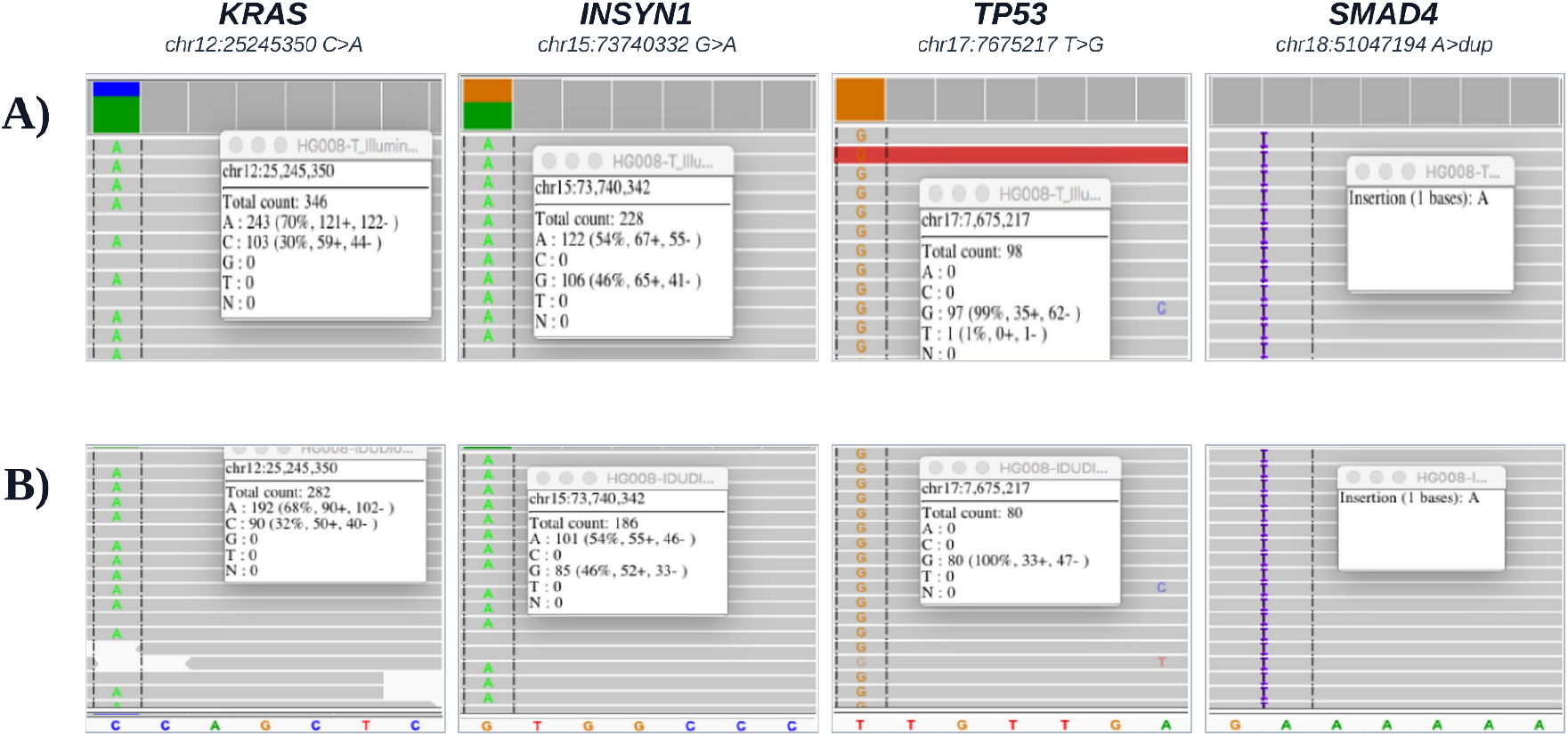
IGV visualization of selected HG008-T variants in the GIAB-provided and Parabricks-derived tumor alignments. Representative loci in *KRAS*, *INSYN1*, *TP53*, and *SMAD4* are shown. (A) GIAB-provided alignment generated with BWA-MEM. (B) Parabricks-derived alignment generated with fq2bam. IGV nucleotide-count pop-ups show the total read depth and allele-specific read counts at each locus.

For the SNVs, IGV variant allele fractions were highly similar between pipelines, with *KRAS* c.35G>T (chr12:25245350 C>A) showing approximately 70% in in the GIAB-provided BWA-MEM versus 68% in Parabricks, *INSYN1* c.457C>T (chr15:73740332 G>A) showing approximately 54% in both alignments, and *TP53* c.395A>C (chr17:7675217 T>G) showing near-complete alternate-allele support in both datasets (approximately 99% in the GIAB-provided BWA-MEM and 100% in Parabricks). Likewise, the *SMAD4* c.153dup locus (chr18:51047194 A>dup) was consistently visualized in both alignments as a 1-bp insertion event.

### BENCHMARKING AGAINST A DRAGEN-DERIVED BASELINE SET USING RTG TOOLS

Haplotype-aware benchmarking with RTG Tools version 3.12.1 of the PASS-filtered, positive-TLOD GPU-derived call set against the PASS-filtered GIAB DRAGEN baseline yielded 14,810 baseline-side true positives and 14,618 call-side true positives before score thresholding, with 2,837 false positives and 2,072 false negatives **(Table 3)**. The positive-TLOD criterion used to construct the input call set was distinct from the subsequently optimized score threshold. Score-based filtering identified 17.520 as the F-measure-maximizing threshold. At this operating point, calls classified as false positives relative to the DRAGEN comparator decreased from 2,837 to 1,392, whereas those classified as false negatives increased from 2,072 to 2,447. This threshold improved precision from 0.8375 to 0.9110 and increased the F-measure from 0.8569 to 0.8821, with a modest reduction in sensitivity from 0.8773 to 0.8551.

**Table 3.** RTG Tools score-based benchmarking summary for the PASS-filtered, positive-TLOD GPU-derived somatic call set against the PASS-filtered GIAB DRAGEN comparator. True-positive, false-positive, and false-negative counts and derived metrics are relative to this comparator, not to an independently validated truth set. The thresholded operating point corresponds to the F-measure-maximizing score threshold.

| **Metric** | **Score threshold = 17.520** | **No score threshold** |
|---|---|---|
| TP (baseline-side) | 14,435 | 14,810 |
| TP (call-side) | 14,245 | 14,618 |
| False positives | 1,392 | 2,837 |
| False negatives | 2,447 | 2,072 |
| Precision | 0.9110 | 0.8375 |
| Sensitivity | 0.8551 | 0.8773 |
| F-measure | 0.8821 | 0.8569 |

The score-dependent behavior of the benchmark is illustrated in **Figure 2**, which shows the RTG ROC-style curve generated from the INFO.RTG_SCORE field. The curve rises steeply at lower false-positive counts and then approaches a plateau, indicating that many baseline-supported variants were recovered before additional false positives accumulated.

**Figure 2.**
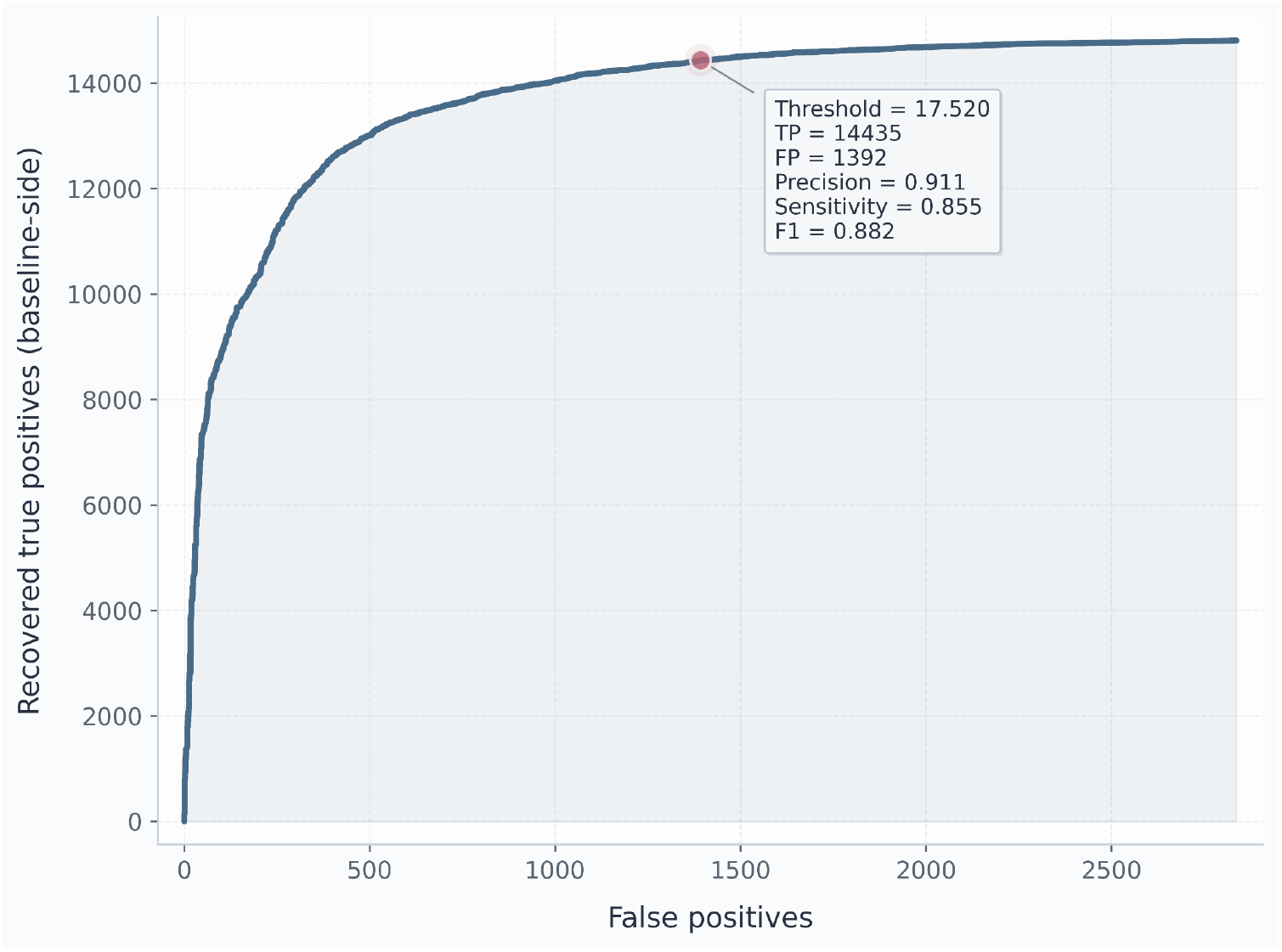
RTG score-based ROC-style curve for the GPU-derived somatic call set compared with the GIAB DRAGEN comparator. The curve was generated from INFO.RTG_SCORE and shows the relationship between false positives and recovered true positives across score thresholds. These classifications are relative to the DRAGEN comparator rather than an independently validated truth set.

### COMPARISON OF SUMMED PREPROCESSING AND VARIANT-CALLING RUNTIMES

Using the fixed CPU–GPU comparison configuration specified in the Methods section **(Table 1)**, the summed runtime of the evaluated preprocessing and somatic variant-calling components was 219 h 24 min 27 s for the CPU-based workflow and approximately 10 h 38 min 32 s for the GPU-accelerated workflow, corresponding to a 20.6-fold speedup **(Figure 3**; **Table 4)**. In the GPU-accelerated workflow, fq2bam performed alignment, sorting, duplicate marking, BQSR table generation, and indexing. BQSR was subsequently applied using separate ApplyBQSR runs.

**Figure 3.**
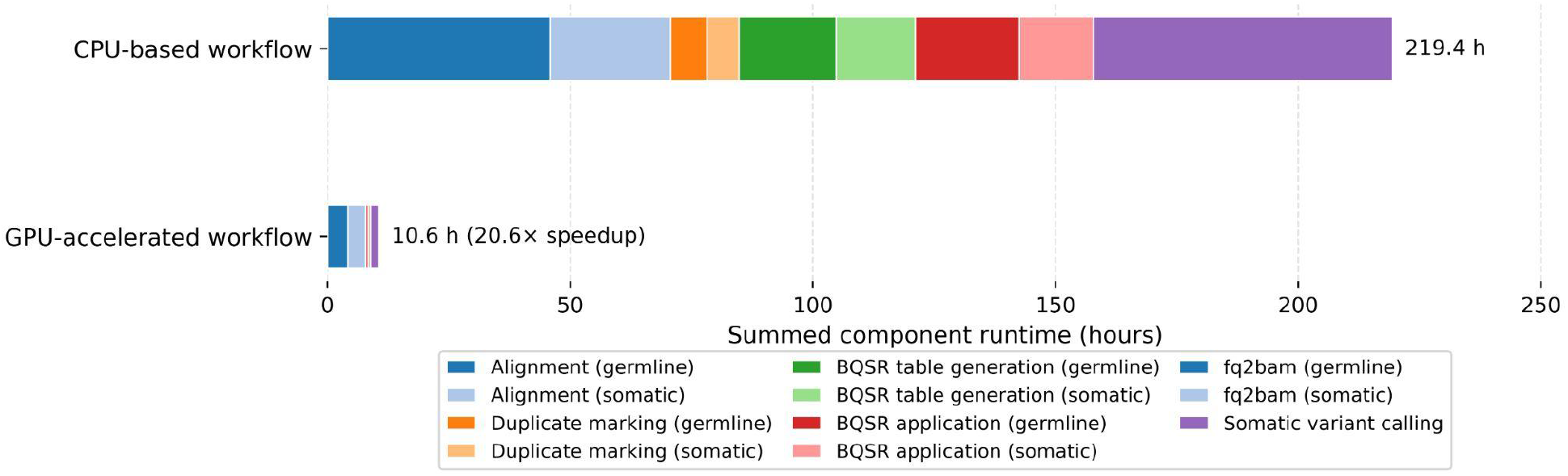
Summed preprocessing and somatic variant-calling runtimes for the CPU-based and GPU-accelerated workflows. CPU preprocessing components are shown separately for germline and somatic data. GPU preprocessing comprises fq2bam and separate ApplyBQSR runs for both samples. The summed runtime was 219 h 24 min 27 s for the CPU-based workflow and approximately 10 h 38 min 32 s for the GPU-accelerated workflow, corresponding to a 20.6-fold speedup.

**Table 4.**
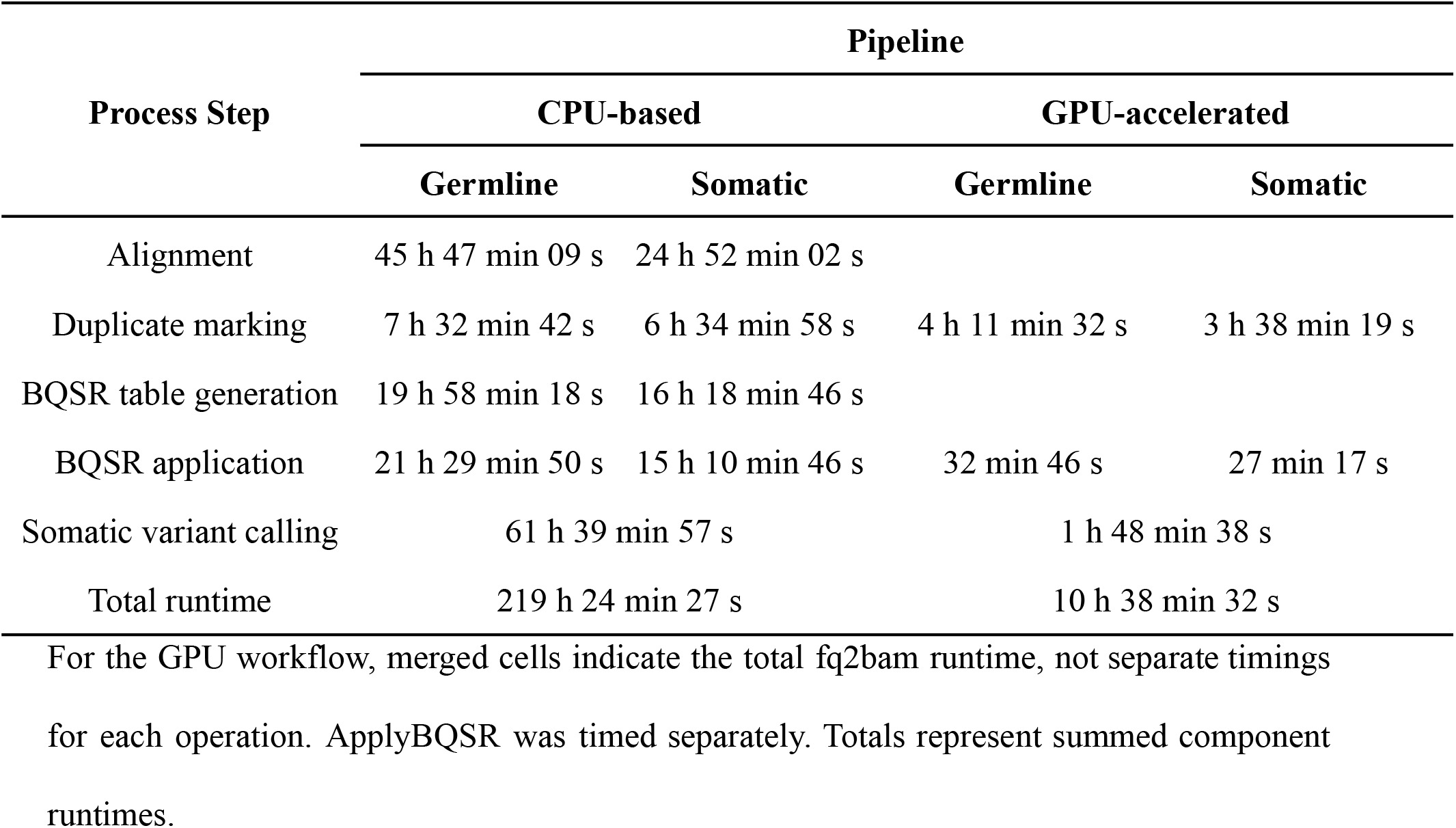
Step-level runtime comparison between the conventional CPU-based workflow and the GPU-accelerated workflow in a tumor–normal design.

| Process Step | Pipeline |  |  |  |
|---|---|---|---|---|
|  | CPU-based |  | GPU-accelerated |  |
|  | Germline | Somatic | Germline | Somatic |
|---|---|---|---|---|
| Alignment | 45 h 47 min 09 s | 24 h 52 min 02 s | 4 h 11 min 32 s | 3 h 38 min 19 s |
| Duplicate marking | 7 h 32 min 42 s | 6 h 34 min 58 s |  |  |
| BQSR table generation | 19 h 58 min 18 s | 16 h 18 min 46 s |  |  |
| BQSR application | 21 h 29 min 50 s | 15 h 10 min 46 s | 32 min 46 s | 27 min 17 s |
| Somatic variant calling | 61 h 39 min 57 s |  | 1 h 48 min 38 s |  |
| Total runtime | 219 h 24 min 27 s |  | 10 h 38 min 32 s |  |

For the GPU workflow, merged cells indicate the total fq2bam runtime, not separate timings for each operation. ApplyBQSR was timed separately. Totals represent summed component runtimes.

In the CPU-based workflow, the summed runtime for germline and somatic preprocessing, including alignment, duplicate marking, BQSR table generation, and BQSR application, was 157 h 44 min 31 s. In the GPU-accelerated workflow, the two fq2bam runs and the two separate ApplyBQSR runs totaled approximately 8 h 49 min 54 s, corresponding to a 17.9-fold speedup.

Somatic variant calling required 61 h 39 min 57 s in the CPU-based workflow using GATK Mutect2 version 4.6.2.0, compared with 1 h 48 min 38 s in the GPU-accelerated workflow using Parabricks MutectCaller version 4.5.1-1, corresponding to an approximate 34.1-fold reduction.

### GPU SCALING AND RUNTIME PERFORMANCE

Runtime profiling showed that the optimal number of GPUs differed across workflow steps and CPU-thread configurations **(Figure 4)**. Under the 16-thread configuration, the fastest somatic alignment runtime was observed with 8 GPUs (00:46:53), whereas germline alignment reached its minimum runtime with 7 GPUs (00:55:22). For variant calling, MutectCaller was fastest with 3 GPUs (01:25:11), while DeepSomatic achieved its shortest runtime with 6 GPUs (01:09:07) **(Figure 4A)**.

**Figure 4.**
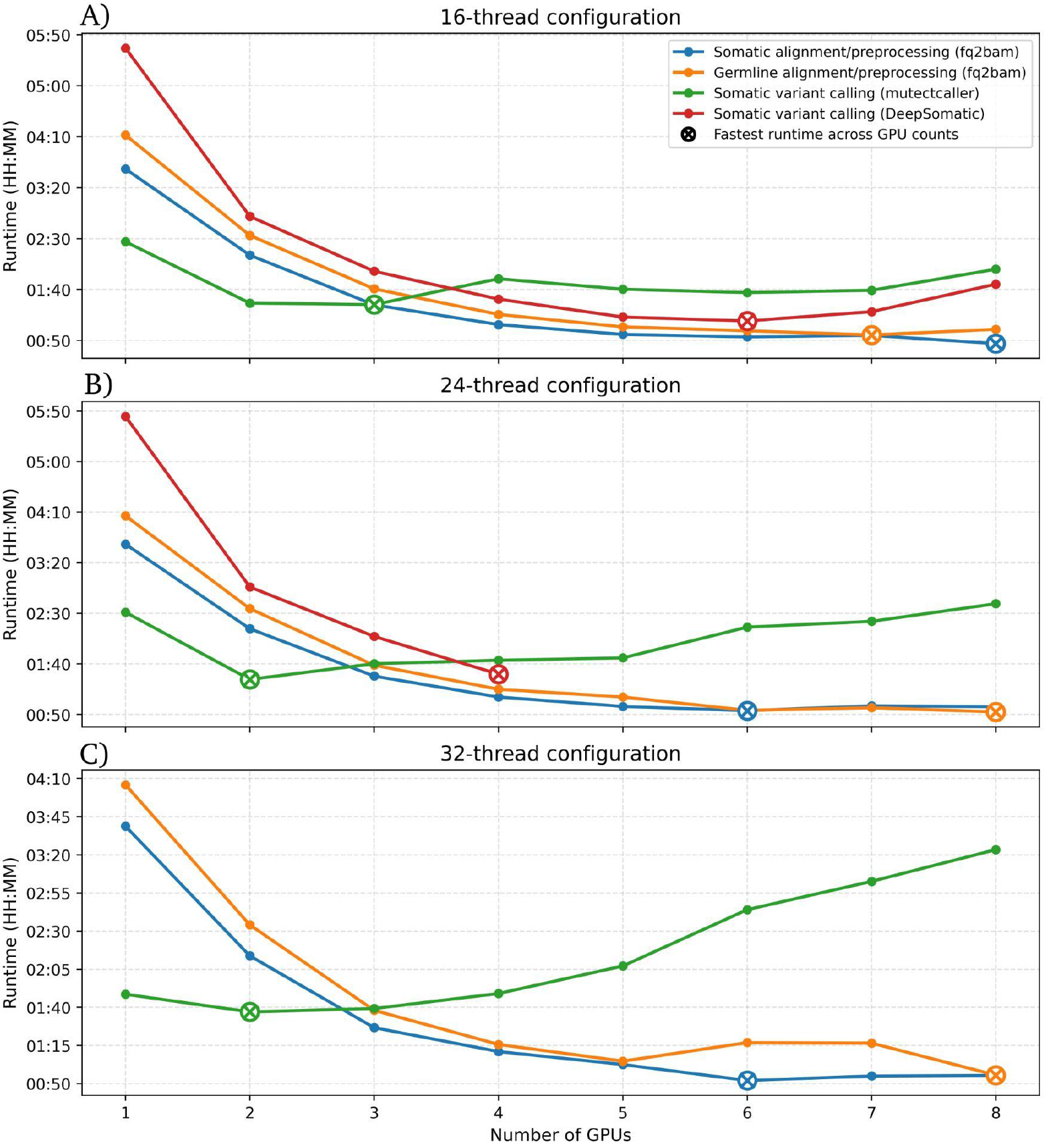
GPU scaling and runtime performance of the Parabricks workflow. Wall-clock runtime across GPU counts for alignment/preprocessing and variant-calling steps under different CPU-thread configurations: (A) 16-thread configuration, (B) 24-thread configuration, and (C) 32-thread configuration. DeepSomatic results under the 32-thread configuration were excluded from formal interpretation because the expected effective CPU-thread allocation exceeded the available host CPU capacity of the DGX A100 server. Complete raw runtime values are provided in **Supplementary Table S2**.

Under the 24-thread configuration, somatic alignment was fastest with 6 GPUs (00:53:46), while germline alignment reached its minimum runtime with 8 GPUs (00:52:31). In contrast, MutectCaller achieved its shortest runtime with only 2 GPUs (01:24:39), after which runtime increased at higher GPU counts. DeepSomatic was evaluated only up to 4 GPUs, where it reached its shortest observed runtime (01:29:52); configurations using more than 4 GPUs were excluded from formal interpretation because the expected effective CPU-thread allocation exceeded the available host CPU capacity of the DGX A100 server **(Figure 4B)**.

Under the 32-thread configuration, somatic alignment again reached its minimum runtime with 6 GPUs (00:52:03), whereas germline alignment was fastest with 8 GPUs (00:55:31). For MutectCaller, the optimal runtime was observed with 2 GPUs (01:37:01), with longer runtimes at higher GPU counts. DeepSomatic results under the 32-thread configuration were excluded from formal interpretation because the expected effective CPU-thread allocation exceeded the available host CPU capacity of the DGX A100 server; therefore, DeepSomatic is not shown in **Figure 4C**.

The fastest CPU-thread/GPU combination differed across workflow steps **(Figure 5)**. Somatic alignment/preprocessing achieved its shortest runtime with 16 threads and 8 GPUs (00:46:53), whereas germline alignment/preprocessing was fastest with 24 threads and 8 GPUs (00:52:31). For MutectCaller, the best overall runtime was obtained with 24 threads and 2 GPUs (01:24:39), although the 16-thread/3-GPU configuration showed a very similar displayed runtime in the heatmap. Among formally interpretable DeepSomatic runs, the fastest runtime was observed with 16 threads and 6 GPUs (01:09:07).

**Figure 5.**
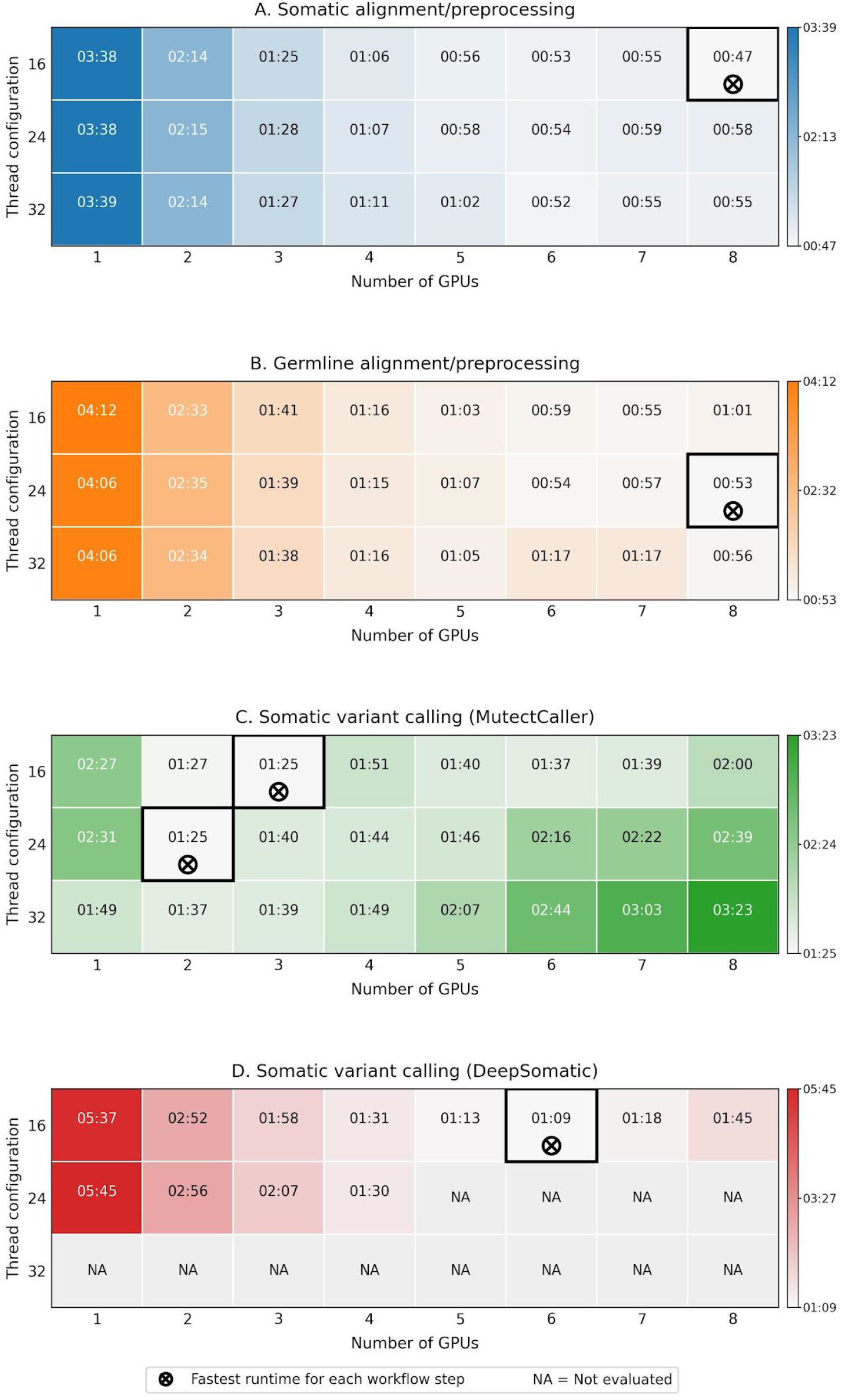
Runtime landscape across GPU and CPU-thread configurations. Heatmaps show wall-clock runtime for (A) somatic alignment/preprocessing, (B) germline alignment/preprocessing, (C) MutectCaller, and (D) DeepSomatic across tested GPU counts and CPU-thread settings. Cell values are shown as HH:MM. The black outlined cell with the circle-and-cross marker indicates the fastest runtime in each panel. NA indicates configurations not evaluated.

### NEXTFLOW EXECUTION AND OUTPUT CONCORDANCE

The full-input HG008 workflow completed all 18 tasks successfully, with no reuse of cached task outputs. The final VCF contained 17,455 records, all with nonempty INFO/CSQ annotations and non-missing positive INFO/TLOD values. Comparison with the VCF generated outside Nextflow identified 17,455 shared CHROM/POS/REF/ALT records and no records exclusive to either output, corresponding to 100% agreement for the compared variant identifiers. The combined task runtime for tumor and normal alignment/preprocessing with fq2bam and somatic variant calling with MutectCaller was 4 h 12 min 20 s.

## DISCUSSION

We evaluated a GPU-accelerated tumor–normal somatic variant-calling pipeline based on NVIDIA Parabricks using the Cancer Genome in a Bottle HG008 dataset. In the fixed comparison, summed preprocessing and somatic variant-calling runtimes were approximately 10 h 38 min 32 s for the GPU-accelerated workflow and 219 h 24 min 27 s for the CPU-based workflow, corresponding to a 20.6-fold speedup. At the analyzed call-set level, intersection-over-union overlap between the PASS-filtered, positive-TLOD Parabricks set and the PASS-filtered DRAGEN comparator was 76.09% for single-nucleotide variants and 69.06% for insertions and deletions.

HG008 was appropriate for this evaluation because it provides publicly available matched tumor and normal WGS data from a pancreatic ductal adenocarcinoma model (20), allowing assessment of GPU-accelerated somatic variant calling in a tumor–normal rather than germline-only benchmark setting.

Variant-level comparisons showed a 73.66% overall overlap between the Parabricks pipeline and the GIAB-provided DRAGEN comparator call set, with higher concordance for SNVs than for indels. Differences between the two call sets were expected because somatic variant detection is influenced by alignment behavior, local sequence context, filtering strategies, and caller-specific modeling assumptions (44). Tumor purity and allelic fraction further influence somatic variant detection sensitivity, particularly for low-frequency variants (45). Consistent with this expectation, low-VAF SNVs were more frequent among pipeline-specific calls than among shared calls. However, more than three quarters of the pipeline-specific SNVs had VAFs of at least 5%, indicating that low VAF alone did not account for most of the observed disagreement. Because 5% was used only as a descriptive threshold and not as a common post hoc filter, this pattern indicates an association between low VAF and discordance rather than exclusion caused by a fixed cutoff. This SNV–indel difference is also consistent with previous benchmarking studies, since indels are generally more sensitive to variant representation and complex genomic contexts than SNVs (46,47).

Haplotype-aware benchmarking with RTG (28) was used to account for representation-related differences between call sets. In this analysis, score-based filtering improved precision from 0.8375 to 0.9110 and F-measure from 0.8569 to 0.8821, indicating that part of the observed disagreement was sensitive to score thresholding rather than fixed across operating points.

Benchmarking guidance emphasizes variant representation, standardized metrics, and stratification by variant type and genomic context. Krusche et al. (2019) also showed that performance can differ markedly inside and outside high-confidence regions (48).

At the time of this analysis, an official high-confidence somatic benchmark BED file was not available for HG008. We therefore compared the whole-genome call sets using DRAGEN as the comparator, without constructing custom confident regions that would require additional methodological assumptions. Germline GIAB benchmarks were not used because they do not directly address tumor–normal somatic variant detection (49). The resulting metrics describe agreement with the DRAGEN call set rather than independently validated analytical accuracy.

The runtime analyses separated the fixed CPU–GPU comparison from the GPU-scaling experiments. For the evaluated components, the GPU-accelerated workflow achieved a 20.6-fold speedup relative to the CPU-based workflow. Summed preprocessing runtime decreased by approximately 17.9-fold, and somatic variant-calling runtime decreased by approximately 34.1-fold.

In the GPU-scaling experiments, NVIDIA Parabricks fq2bam tool and MutectCaller showed different scaling patterns. Alignment and preprocessing with NVIDIA Parabricks fq2bam tool benefited from increasing GPU allocation and showed comparatively limited sensitivity to the tested CPU-thread settings. This pattern is consistent with NVIDIA Parabricks fq2bam tool integration of BWA-MEM-based alignment, sorting, duplicate marking, and BQSR report generation within a GPU-accelerated workflow (38). Under the tested configurations, fq2bam scaling was more responsive to GPU count than to the evaluated CPU-thread settings, although CPU resources may still contribute to preprocessing, data transfer, and execution overhead.

In contrast, MutectCaller was fastest with 2–3 GPUs, and higher GPU counts did not improve runtime under the tested configurations. For example, under the 24-thread configuration, MutectCaller achieved its shortest runtime with 2 GPUs (01:24:39), whereas higher GPU counts resulted in progressively longer runtimes. This pattern is consistent with previous Parabricks benchmarking by O’Connell et al. (2023), who reported strong acceleration across genomic workflows overall but more variable scaling for somatic callers across GPU counts and computing platforms (18). In our study, the longer runtimes observed at higher GPU counts indicate that additional GPUs can introduce coordination or data-movement overhead for MutectCaller under these conditions.

DeepSomatic was included as a deep-learning–based somatic small-variant caller for comparison with MutectCaller (50). In our experiments, DeepSomatic benefited from multi-GPU execution under feasible configurations, but scaling was constrained by host-thread demand as GPU count and stream allocation increased. Under the tested conditions, DeepSomatic configurations should be selected according to stream count, GPU count, and the resulting host-thread demand.

MutectCaller and DeepSomatic runtimes are not directly comparable because the tested implementations used different preprocessing inputs. MutectCaller used reheadered alignment files together with recalibration tables generated during preprocessing, whereas DeepSomatic was executed directly from tumor and normal alignment files without incorporating base quality score recalibration tables. Consequently, observed differences likely reflect both algorithmic characteristics and differences in preprocessing requirements rather than intrinsic computational efficiency alone.

The entire evaluation was based on matched tumor–normal WGS data from a single individual, HG008. Although HG008 provides a well-characterized, publicly available somatic WGS model, additional evaluations across tumor types, sequencing depths, tumor purities, and somatic variant allele-fraction distributions are needed to assess the generalizability of the observed concordance and computational performance. The runtime experiments measured processing time for this tumor–normal pair rather than throughput under concurrent processing of multiple samples, which may vary with workload and resource allocation (51).

Across the scaling experiments, the fastest configurations differed by workflow step: NVIDIA Parabricks fq2bam tool performed best with 6–8 GPUs, whereas MutectCaller reached its shortest runtime with 2–3 GPUs. These results favor step-specific tuning of accelerated somatic workflows over uniform allocation of the maximum available GPU count.

## CONCLUSION

In this HG008 tumor–normal WGS analysis, the GPU-accelerated workflow achieved a 20.6-fold speedup in summed preprocessing and somatic variant-calling runtime relative to the CPU-based workflow. GPU-scaling experiments showed that the fastest configurations differed between preprocessing and somatic variant calling. With score-based filtering, haplotype-aware RTG benchmarking against the DRAGEN-derived comparator call set increased the F-measure from 0.8569 to 0.8821; IGV manual review also showed consistent read-level support at the selected HG008-T loci. GPU resources should therefore be assigned by workflow step rather than maximized uniformly across the pipeline, and variant-level performance should be reported as comparator-based benchmarking rather than truth-set accuracy. The Nextflow implementation also reproduced the historical set of 17,455 CHROM/POS/REF/ALT records using all HG008 reads.

## List of Abbreviations

BAM: Binary Alignment Map
BCFtools: Binary Variant Call Format tools.
BQSR: Base Quality Score Recalibration.
CPU: Central Processing Unit.
DRAGEN: Dynamic Read Analysis for GENomics.
FASTQ: FASTQ sequence format.
GIAB: Genome in a Bottle.
GPU: Graphics Processing Unit.
GRCh38: Genome Reference Consortium Human Build 38.
IGV: Integrative Genomics Viewer.
Indels: Insertions and deletions
MD5: Message Digest Algorithm 5
NGS: Next Generation Sequencing.
PASS: Variants passing filtering criteria
ROC: Receiver Operating Characteristic
RTG: Real Time Genomics.
SDF: Sequence Data Format
SNVs: Single Nucleotide Variants
TLOD: Tumor Log Odds score.
VAF: Variant Allele Fraction.
VCF: Variant Call Format.
WGS: Whole Genome Sequencing.

## Declarations

### Ethics approval and consent to participate

Not applicable

### Consent for publication

Not applicable.

### Availability of data and materials

The sequencing data and comparator call sets used in this study are publicly available from the Genome in a Bottle data repository, including the HG008 sequencing data (21), the DRAGEN-derived comparator call set (22), and the corresponding checksum files (23). The BRISANCE repository contains the pipeline source code, historical workflows, variant-set comparison and RTG benchmarking scripts, Nextflow implementation, and user configuration interface (26). Installation and usage documentation, including video tutorials, are available on the BRISANCE website (52).

### Competing interests

The authors declare no competing interests.

### Funding

This work was supported by the Conselho Nacional de Desenvolvimento Científico e Tecnológico (CNPq) through Chamada n° 16/2023 – Saúde de Precisão, as part of the InovaCriança project, Inovação terapêutica em crianças com tumores sólidos: uma jornada de precisão — project MS-SECTICS-Decit/CNPq n° 444217/2023-1 — funded by the Department of Science and Technology of the Secretariat of Science, Technology, Innovation and Strategic Inputs of the Brazilian Ministry of Health. MPA is a recipient of a CAPES doctoral fellowship. RMR is a recipient of a CNPq Productivity fellowship.

### Authors’ contributions

**MPA:** Data curation; Formal analysis; Investigation; Methodology; Software; Visualization; Writing – original draft; Project administration. **SB:** Writing – review & editing. **SAT:** Funding acquisition; Writing – review & editing. **RMR:** Supervision; Funding acquisition; Project administration; Writing – review & editing. **LSS:** Conceptualization; Writing – review & editing. **LGCS:** Supervision; Project administration; Writing – review & editing. All authors read and approved the final manuscript.

### Artificial Intelligence Disclaimer

Language revision was performed using Grammarly (Grammarly Inc., USA; version 1.2.283.1934) for grammar, spelling, punctuation, clarity, and stylistic consistency. Codex (OpenAI) was used to assist with code review and deployment through GitHub and Vercel. The authors declare that no generative artificial intelligence tools were used to write, generate, or substantively modify the scientific content of this manuscript.

## Acknowledgments

Computational resources were provided by the Artificial Intelligence Applied to Oncology research group at Barretos Cancer Hospital. The group received support from the Brazilian Ministry of Science, Technology and Innovation through Funding Agreement No. 947017/2023. We thank Dr. André Luiz Pinto Santos and Dr. Vinícius Silva Madureira Pereira for server administration and technical support.

## Supplementary information

Supplementary Material 1 (XLSX). Supplementary Table S1 presents the VAF distribution and cross-pipeline characterization of shared and pipeline-specific SNVs. Supplementary Table S2 presents the complete runtime values across the evaluated CPU-thread and GPU configurations.

## Notes

### Competing Interest Statement

The authors have declared no competing interest.

